# Restrained activation of CYFIP2-containing WAVE complexes controls membrane protrusions and cell migration

**DOI:** 10.1101/2020.07.02.184655

**Authors:** Anna Polesskaya, Arthur Boutillon, Sheng Yang, Yanan Wang, Stéphane Romero, Yijun Liu, Marc Lavielle, Nicolas Molinie, Nathalie Rocques, Artem Fokin, Raphaël Guérois, Baoyu Chen, Nicolas B. David, Alexis M. Gautreau

## Abstract

Branched actin networks polymerized by the Arp2/3 complex are critical for cell migration. The WAVE complex is the machinery that activates Arp2/3 in a RAC1-dependent manner at the leading edge of migrating cells. Multiple WAVE complexes are assembled in a cell through various combinations of paralogous subunits. Here we report the surprising phenotype associated with loss-of-function of CYFIP2, a subunit of the WAVE complex. In three different human mammary cell lines and in prechordal plate cells of gastrulating zebrafish embryos, CYFIP2 depletion promoted, rather than impaired, membrane protrusions and migration persistence. CYFIP2, however, assembled WAVE complexes that polymerize branched actin at the cell cortex and rescued membrane protrusions of *CYFIP1/2* double knock-out cells, although less efficiently than CYFIP1. Point mutations of CYFIP2 associated with intellectual disability in children were gain-of-function, as they made CYFIP2 as active as CYFIP1 in this rescue experiment. Biochemical reconstitutions of CYFIP2-containing WAVE complexes showed that they bound equally well to active RAC1 as CYFIP1-containing WAVE complexes, yet they were poorly activated in response to RAC1 binding. Together these results suggest that CYFIP2-containing WAVE complexes titrate active RAC1 and thereby prevent efficient CYFIP1-containing complexes from being activated. In this context, where cell migration is governed by the balance of CYFIP1/2 expression, releasing the restrained activity of CYFIP2-containing WAVE complexes leads to pathology.

## Introduction

Polymerization of branched actin powers adherent membrane protrusions, known as lamellipodia, which are critical for cell migration. The Arp2/3 complex generates branched actin when activated by Nucleation Promoting Factors (NPFs) (Rotty *et al*, 2013). At the cell cortex, WAVE proteins are the major NPFs that induce lamellipodia. WAVE proteins are regulated within a stable pentameric complex (Rottner *et al*, 2021). The WAVE complex maintains WAVE in an inactive conformation by masking its output Arp2/3 activating VCA domain (Derivery *et al*, 2009; Ismail *et al*, 2009), but at the leading edge, RAC1 signaling unmasks the VCA from the WAVE complex, and thereby activates Arp2/3 (Chen *et al*, 2010; Steffen *et al*, 2013; Schaks *et al*, 2018; Mehidi *et al*, 2019). Two GTP-bound RAC1 molecules bind to the CYFIP subunit to activate the WAVE complex (Chen *et al*, 2017; Ding *et al*, 2022). The RAC1-WAVE-Arp2/3 pathway controls membrane protrusions and for how long this protrusion is sustained (Krause & Gautreau, 2014). A lamellipodium represents a persistent way of migrating. Arpin, an Arp2/3 inhibitory protein, which is also under the control of RAC1, antagonizes the RAC1-WAVE-Arp2/3 pathway, renders the lamellipodium unstable and decreases migration persistence (Dang *et al*, 2013).

The WAVE complex is a combination of subunit isoforms. A WAVE complex is composed of 5 generic subunits, referred to as CYFIP, NAP, WAVE, ABI and BRK1. Except BRK1, all human subunits are encoded by paralogous genes, 2 for CYFIP and NAP, 3 for WAVE and ABI (Derivery and Gautreau, 2010). There are as many as 2×2×3×3, *i.e*. 36, possible WAVE complexes, just by combining the different paralogous subunits. Furthermore, the *ABI1* gene has been shown to be alternatively spliced and the resulting isoforms do not possess the same ability to mediate macropinocytosis, which, like lamellipodium formation, depends on the ability of branched actin to drive membrane protrusions (Dubielecka et al., 2010). In mouse embryonic fibroblasts, WAVE2 is critical for the formation of peripheral ruffles, whereas WAVE1 is critical for dorsal ruffles (Suetsugu et al., 2003). In melanoma cells, however, WAVE1 and WAVE2 are redundant in lamellipodium formation (Tang *et al*, 2020). Thus, evidence exists for some functional specialization among WAVE complexes in an overall context of redundancy.

CYFIP1 and CYFIP2 are co-expressed in many cell types. They are thought to be redundant and were simultaneously knocked out to obtain melanoma cells fully devoid of lamellipodia (Schaks *et al*, 2018). In neurons, the two paralogs might have specific functions. For example, in retinal ganglion cells, CYFIP2 is specifically involved in axon sorting (Pittman *et al*, 2010; Cioni *et al*, 2018). Moreover, the *CYFIP2* gene is specifically mutated in children affected by intellectual disability and epileptic encephalopathy (Nakashima *et al*, 2018; Zweier *et al*, 2019; Begemann *et al*, 2021).

We sought to compare the roles of the two CYFIP paralogs, CYFIP1 and CYFIP2, in cell migration and found a striking apparent anti-migratory function of CYFIP2 in many different cell systems. This anti-migratory function seems at odds with the role of WAVE complexes in activating Arp2/3, but can be accounted for by the specific regulation that CYFIP2 confers to the WAVE complex and the balance between CYFIP1- and CYFIP2-containing complexes in the cell.

## Results

### Depletion of CYFIP2 yields a phenotype at odds with a WAVE complex subunit

We depleted CYFIP1 and CYFIP2 from MDA-MB-231 mammary carcinoma cells using pools of four siRNAs (Smartpools from Dharmacon). As a control, we also depleted NCKAP1, which is an obligate subunit in the composition of WAVE complexes in mammary cells, since its paralog, NCKAP1L, also known as HEM1, is restricted to hematopoietic cells (Derivery & Gautreau, 2010b). Depletion of these different subunits had different impacts on the WAVE complexes expressed by MDA-MB-231 cells. Indeed, WAVE complex subunits are stable when assembled. Depletion of a subunit usually destabilizes the multiprotein complex that contains it (Derivery & Gautreau, 2010b). As expected, depletion of NCKAP1 led to a severe downregulation of WAVE complex subunits, including CYFIP1 and CYFIP2 (Fig.1A). This result suggests that CYFIP1 and CYFIP2 coexist as different WAVE complexes, all of which depending on NCKAP1. Depletion of CYFIP1 led to a significant destabilization of WAVE complexes, as appreciated by levels of NCKAP1 and WAVE2. In contrast, depletion of CYFIP2 did not lead to any visible depletion of the other subunits. We thus decided to measure absolute levels of CYFIP1 and CYFIP2 by Western blots using purified CYFIP1- or CYFIP2-containing WAVE complexes as standards (Fig.S1). We found that MDA-MB-231 cells contain 208 pg of CYFIP1 and 25 pg of CYFIP2 per μg of extract (Fig.1B). The fact that CYFIP1 is 8-fold more abundant than CYFIP2 is in line with the destabilization of WAVE complex subunits that was more obvious when CYFIP1 was depleted than when CYFIP2 was depleted. The depletion of either CYFIP protein also resulted in a compensatory up-regulation of the remaining CYFIP paralogous subunit. In particular, CYFIP2 levels were increased by 46 % when CYFIP1 was depleted (Fig.S1).

**Figure 1.**
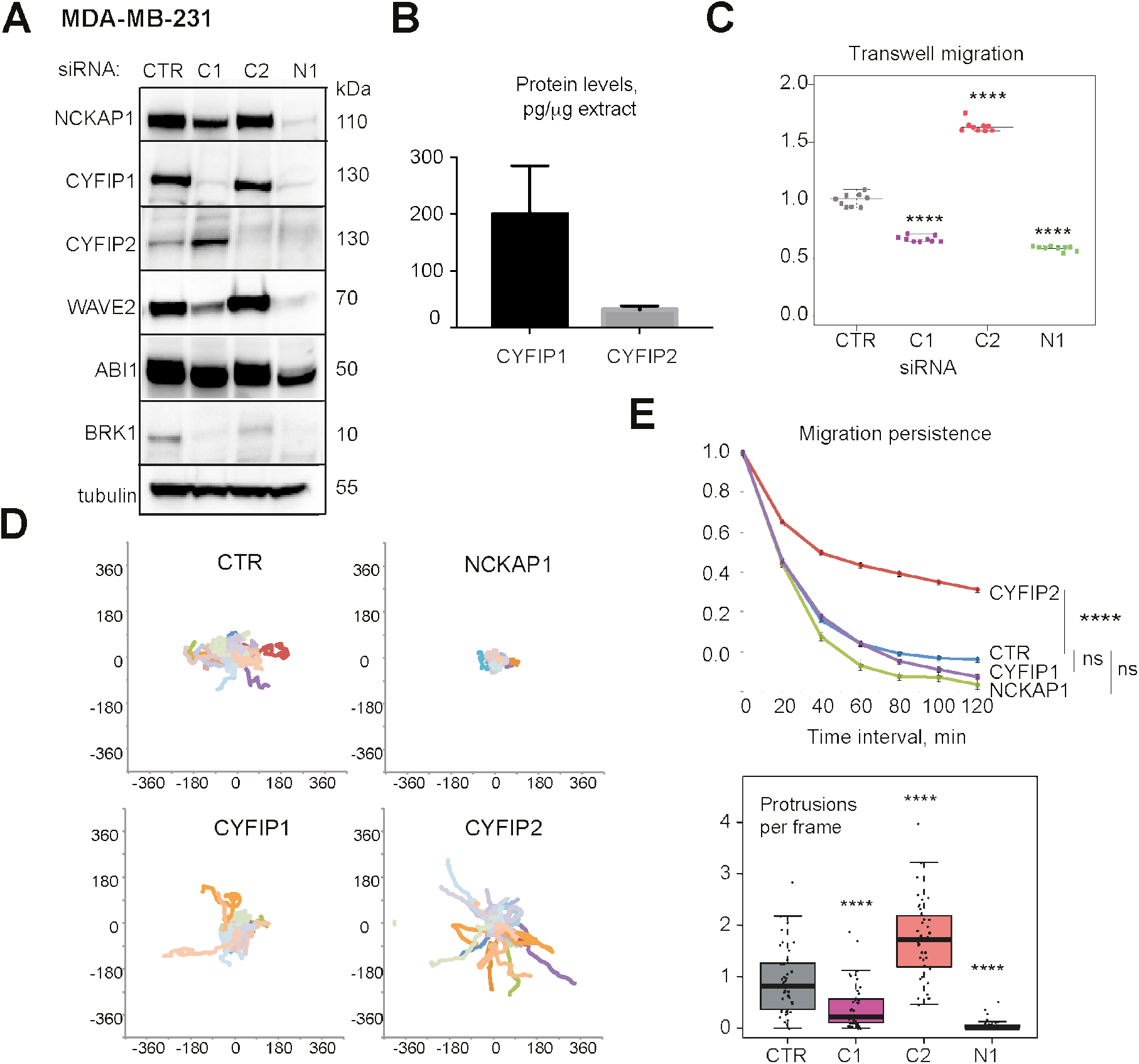
Depletion of the WAVE complex subunit CYFIP2 promotes migration of MDA-MB-231 cells. **A** MDA-MB-231 cells were transfected with pools of siRNAs targeting CYFIP1 (C1), CYFIP2 (C2), NCKAP1 (N1) or non-targeting siRNAs (CTR). Western blots of WAVE complex subunits and tubulin as a loading control. **B** Levels of CYFIP1 and CYFIP2 proteins in RIPA extract of MDA-MB-231 cells. Mean ± SD of 3 biological repeats. **C** Quantification of transwell migration efficiency of transfected MDA-MB-231 cells. Mean ± SD of 9 measurements (3 biological repeats performed in triplicate). **D** MDA-MB-231 cells depleted of the indicated proteins were embedded into 3D collagen type I gels and their trajectory recorded. **E** Migration persistence of 3D MDA-MB-231 cells and average number of protrusions per frame are plotted. Mean ± sem. Box and whiskers plot, n=30. One-Way ANOVA, * P<0.05; ** P<0.01; *** P<0.001; **** P<0.0001; ns not significant.

We first used a transwell assay to evaluate the migration of MDA-MB-231 cells when CYFIP1, CYFIP2, or NCKAP1 was depleted. As expected, depletion of NCKAP1 and CYFIP1 significantly decreased the number of cells that migrated through the filter (Fig.1C). To our surprise, depletion of CYFIP2 had the inverse effect, since it significantly promoted migration through the filter. We then attempted to confirm this intriguing result with stable depletion using shRNA. We derived MDA-MB-231 lines expressing either a shRNA targeting NCKAP1 or CYFIP2 using published plasmids (Steffen *et al*, 2004). We also obtained stable MDA-MB-231 lines overexpressing CYFIP2, but we were unable to obtain clones expressing NCKAP1 in parallel selection schemes. shRNA-mediated depletion of CYFIP2 increased cell migration in the transwell assay, whereas CYFIP2 overexpression decreased it (Fig.S2). The function of CYFIP2 is thus to inhibit, rather than to promote, cell migration in both loss- and gain-of function experiments.

We then turned to a more physiopathological assay for these invasive carcinoma cells. We seeded isolated MDA-MB-231 cells in 3D collagen gels. In these settings, phenotypic differences between the various siRNA-depleted cells were dramatic for both migration and survival. NCKAP1-depleted cells died during the first 24 h (Movie S1). CYFIP1-depleted cells also appeared prone to die in these conditions. Before they died, these cells migrated and we compared their trajectories to control cells or CYFIP2-depleted cells, which did not show any sign of cell death (Fig.1D). In this assay, CYFIP2-depleted cells formed more protrusions and migrated more persistently than the other cells they were compared with (Fig.1E). Other migration parameters, such as cell speed and Mean Square Displacement (MSD) were only marginally affected in CYFIP2-depleted cells, in contrast to NCKAP1-depleted cells, where all migration parameters were dramatically decreased (Fig.S3). Because in 3D collagen gels, cell death was present in addition to migration, we further evaluated the surprising role of CYFIP2 in other cell migration assays.

In a wound healing assay, CYFIP2 depletion also promoted migration of MDA-MB-231 cells unlike the depletion of NCKAP1 and CYFIP1 (Fig.S2). We sought to confirm the MDA-MB-231 result with MCF7 cells, where wound healing corresponds to collective migration, since MCF7 cells are epithelial, whereas MDA-MB-231 cells are mesenchymal. When we depleted NCKAP1, CYFIP1 and CYFIP2 from MCF7 cells, we observed by Western blot a similar effect on subunit stability as in MDA-MB-231 cells (Fig.2A). CYFIP2 depletion promoted collective migration, since these cells healed the wound in 20 h compared with 30 h for control cells (Movie S2). In contrast, CYFIP1- or NCKAP1-depleted cells were dramatically less efficient at closing the wound – it took them 78 and 81 h respectively – as expected for cells deficient in WAVE complexes (Fig. 2B).

**Figure 2.**
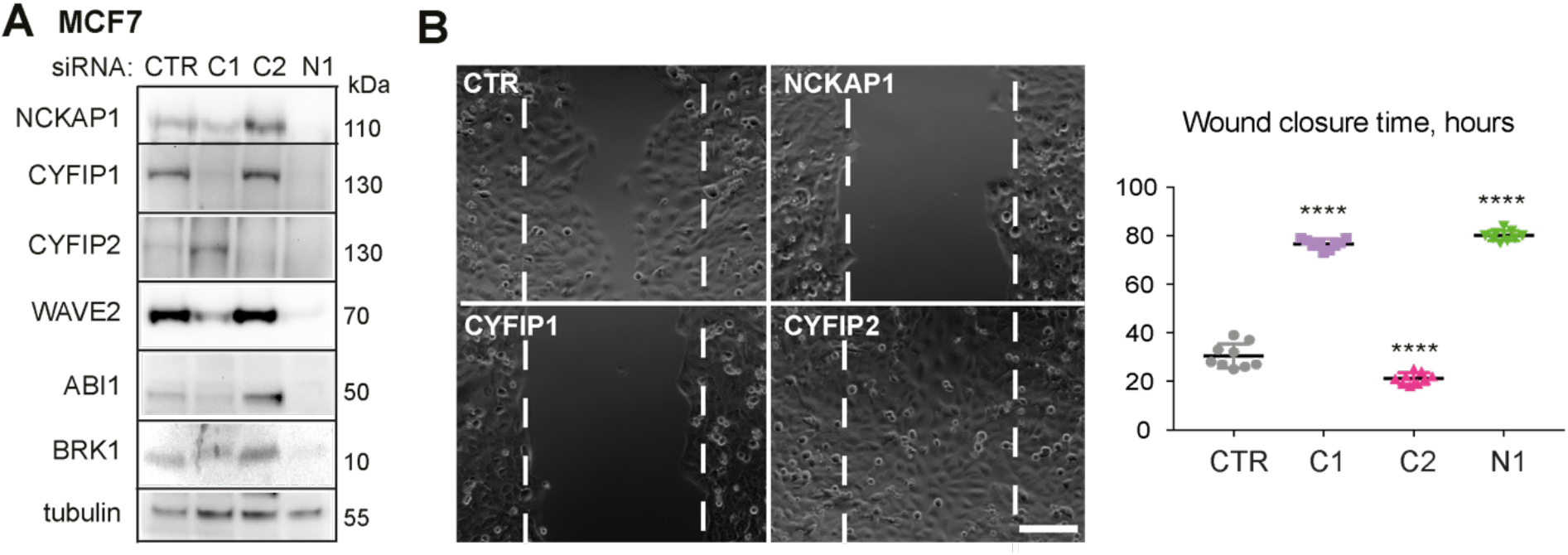
CYFIP2 depletion promotes wound healing of MCF7 cells. **A** MCF7 cells were transfected with pools of siRNAs targeting CYFIP1 (C1), CYFIP2 (C2), NCKAP1 (N1) or nontargeting siRNAs (CTR). Western blots of WAVE complex subunits and tubulin as a loading control. **B** Wound healing of MCF7 cells. Still phase contrast images corresponding to the time, when the first wound is healed (CYFIP2) and plots corresponding to the time taken to close the wound. Mean ± SD of 9 measurements (3 biological repeats performed in triplicate). One-Way ANOVA, **** P<0.0001. Scale bar: 400 μm.

In an attempt to further generalize these results, we turned to the immortalized, but not transformed, MCF10A mammary cell line. MCF10A cells contain 239 pg of CYFIP1 and 39 pg of CYFIP2 per μg of extract (Fig.3A). MCF10A cells thus express approximately 6-fold more CYFIP1 than CYFIP2. Depletions of NCKAP1, CYFIP1 or CYFIP2 using siRNA pools in MCF10A cells were efficient and yielded similar results on the levels of WAVE complex subunits as in MDA-MB-231 and MCF7 cell lines (Fig.3B). The depletion of either CYFIP protein also resulted in a compensatory up-regulation of the remaining CYFIP paralogous subunit. In particular, CYFIP2 levels were increased by 167 % when CYFIP1 was depleted (Fig.S1).

**Figure 3.**
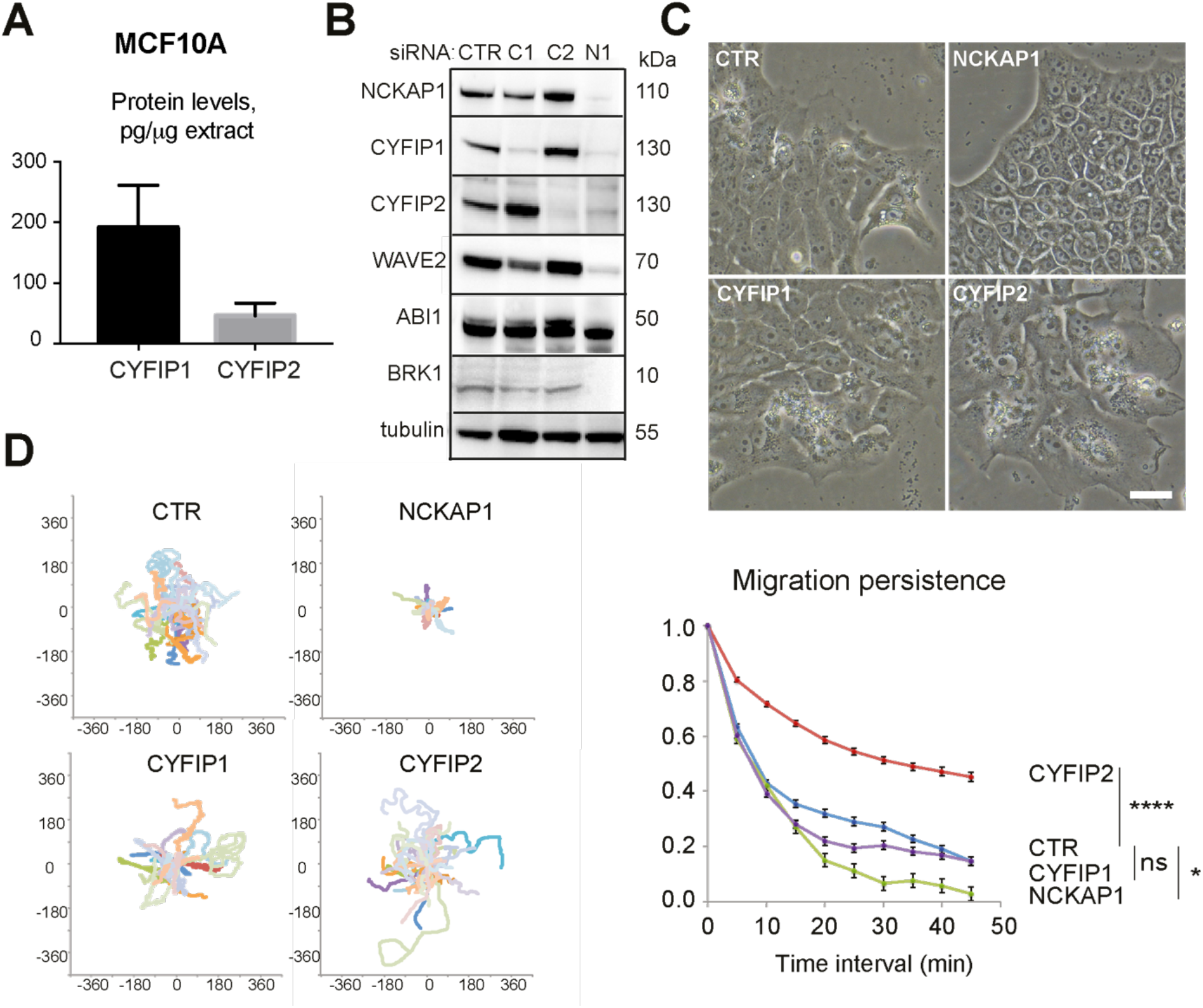
CYFIP2 depletion promotes persistent migration of MCF10A cells. **A** Levels of CYFIP1 and CYFIP2 proteins in MCF10A cells. Mean ± SD of 3 biological repeats. **B** MCF10A cells were transfected with pools of CYFIP1 (C1), CYFIP2 (C2), NCKAP1 (N1) or non-targeting siRNAs (CTR). WAVE complex subunits and tubulin as a loading control were analyzed by Western blot. **C** Phase-contrast images of depleted cells. Scale bar: 50 μm. **D** Trajectories of single MCF10A cells migrating on fibronectin-coated 2D substrate and corresponding migration persistence, n=25 cells. One-way ANOVA, * P<0.05; **** P<0.0001; ns not significant.

MCF10A cells establish cell-cell junctions and form epithelial islets. However, MCF10A cells also frequently migrate as single cells in 2D cultures in their regular medium containing EGF. In these conditions, NCKAP1-depleted cells appeared tightly packed in epithelial islets, whereas CYFIP2-depleted cells appeared on the opposite, more spread with large membrane protrusions, even if cells remained associated with one another (Fig.3C, Movie S3). CYFIP1 depletion did not have a pronounced effect on cell morphology. We then recorded siRNA-treated MCF10A cells to analyze cell migration. Trajectories of single cells were plotted (Fig.3D). Again, we observed that depletion of CYFIP2 dramatically increased migration persistence, unlike depletion of CYFIP1 or NCKAP1. Cell speed and MSD were only marginally affected (Fig.S3). These phenotypes obtained with pools of 4 siRNAs were confirmed with 2 single siRNA sequences (Fig.S4).

We further validated our results using *CYFIP2* knock-out (KO) clones created by CRISPR-Cas9 in MCF10A cells. We isolated two CYFIP2 negative clones, where both alleles were inactivated due to insertions/deletions producing a frameshift (Fig.S5). Both *CYFIP2* KO clones displayed increased migration persistence in single cell migration assays, as in siRNA-depleted MCF10A cells (Fig.S5). This CYFIP2 loss-of-function phenotype with a systematic increase in migration persistence was also the phenotype reported when the small GTPase RAC1 was activated by the Q61L mutation, or when the Arp2/3 inhibitory protein Arpin was depleted (Dang *et al*, 2013; Molinie *et al*, 2019). In these various assays using 3 different cell lines, CYFIP2 appears to antagonize the RAC1-WAVE-Arp2/3 pathway, even though it is a subunit of the WAVE complex that activates the Arp2/3 complex.

### Depletion of CYFIP1 and CYFIP2 yield opposite phenotypes in zebrafish embryos

To validate this apparent anti-migratory function of CYFIP2 in a physiological system, we turned to the zebrafish embryo. During gastrulation, prechordal plate cells collectively migrate from the fish organizer (shield) to the animal pole of the embryo by forming actin-rich protrusions (Montero *et al*, 2003; Dumortier *et al*, 2012). These RAC1-dependent protrusions are the 3D equivalents of 2D lamellipodia and are easily distinguished from thin, filopodia-like extensions (Diz-Muñoz *et al*, 2010; Petrie *et al*, 2012). We assessed the function of endogenous CYFIP1, CYFIP2 and NCKAP1 using morpholino-mediated loss-of-function and rescue using injection of mRNAs encoding the corresponding human subunits (Fig.4A).

**Figure 4.**
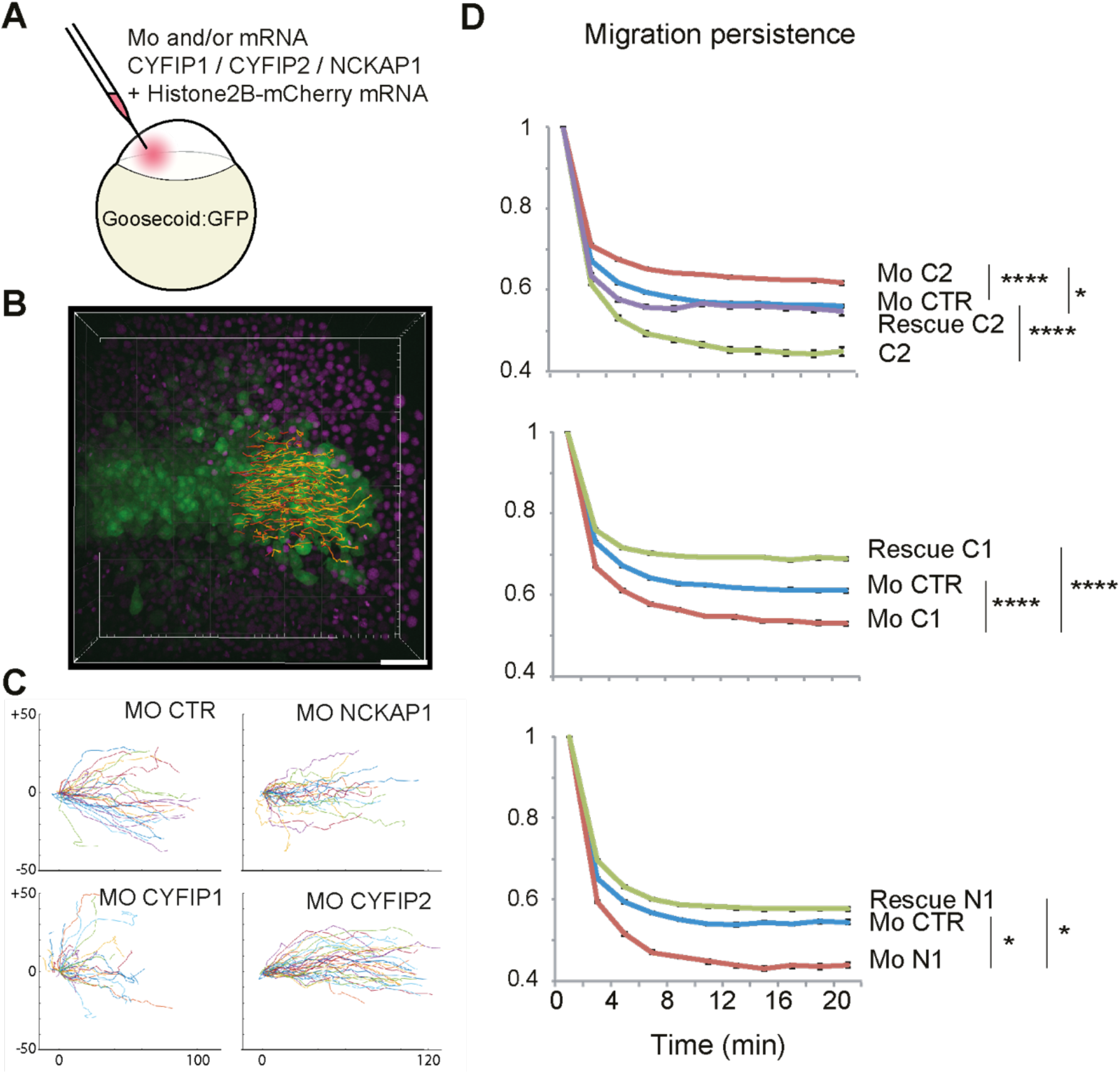
CYFIP2 depletion promotes migration persistence of prechordal plate cells during gastrulation of zebrafish embryos. **A** Embryos were injected with Histone2B-mCherry mRNA and morpholinos (Mo) targeting a control sequence (CTR), CYFIP1 (C1), CYFIP2 (C2), NCKAP1 (N1), alone or in combination with mRNAs encoding the corresponding human protein (rescue). **B** Dorsal view of a volume acquisition of a Tg(Gsc:GFP) zebrafish embryo. Scale bar: 50 μm. Animal pole is located to the right. Notochord and prechordal plate cells express GFP (green) and nuclei express histone2B-mCherry (in magenta). **C** Nuclei of prechordal plate cells were tracked in 3D over time. Trajectories of 50 randomly selected cells (10 first time points, i.e. 20 min) in each condition were plotted from the same origin. Axes are in μm. **D** Migration persistence of prechordal plate cells injected with the indicated Mo and/or mRNA. C2 corresponds to the simple expression of C2, whereas Rescue C2 corresponds to C2 expression together with the morpholino targeting endogenous C2. More than 500 cells (from 4 to 6 embryos pooled from experiments performed on different days, with 100 to 200 cells per embryo) were analyzed per condition. One-Way ANOVA, * P<0.05; **** P<0.0001.

We first analyzed cell trajectories in embryos injected with morpholino and/or mRNA for CYFIP1, CYFIP2 and NCKAP1 (Fig.4A). Experiments were performed in a goosecoid:GFP transgenic line, allowing easy identification of prechordal plate cells. Nuclei were labeled by expression of a Histone2B–mCherry construct for easy cell tracking (Movie S4), and cell trajectories were plotted (Fig.4B). Similar to what was observed in human cell lines, CYFIP2 depletion increased migration persistence of zebrafish prechordal plate cells compared to injection of a control morpholino (Fig.4C). This effect was rescued by co-injection of the morpholino-insensitive human CYFIP2 mRNA (Fig.4D), demonstrating the specificity of the phenotype. Consistently, overexpression of CYFIP2, i.e. injection of the same amount of mRNA as for the rescue but without the corresponding morpholino, decreased cell persistence. In contrast to CYFIP2, downregulation of CYFIP1 or NCKAP1 reduced cell persistence, both effects being rescued by co-injection of their corresponding mRNAs.

We then used cell transplants to examine whether the effect was cell autonomous and to analyze dynamics of cell protrusions. Some prechordal plate cells from a donor embryo injected with morpholino and/or mRNA were transplanted to the prechordal plate of an untreated host embryo (Fig.5A). Actin-rich protrusions were highlighted by the enrichment of the Lifeact-mCherry marker (Fig.5B, Movie S5). CYFIP2 depletion doubled the number of protrusions and increased protrusion length compared to cells injected with a control morpholino (Fig.5C). These two effects were rescued by CYFIP2 mRNA. Consistently, CYFIP2 overexpression decreased the number of protrusions, similar to the effect of depleting NCKAP1 or CYFIP1. CYFIP2 overexpression did not affect protrusion length, but this parameter was also not affected by depletions of NCKAP1 or CYFIP1.

**Figure 5.**
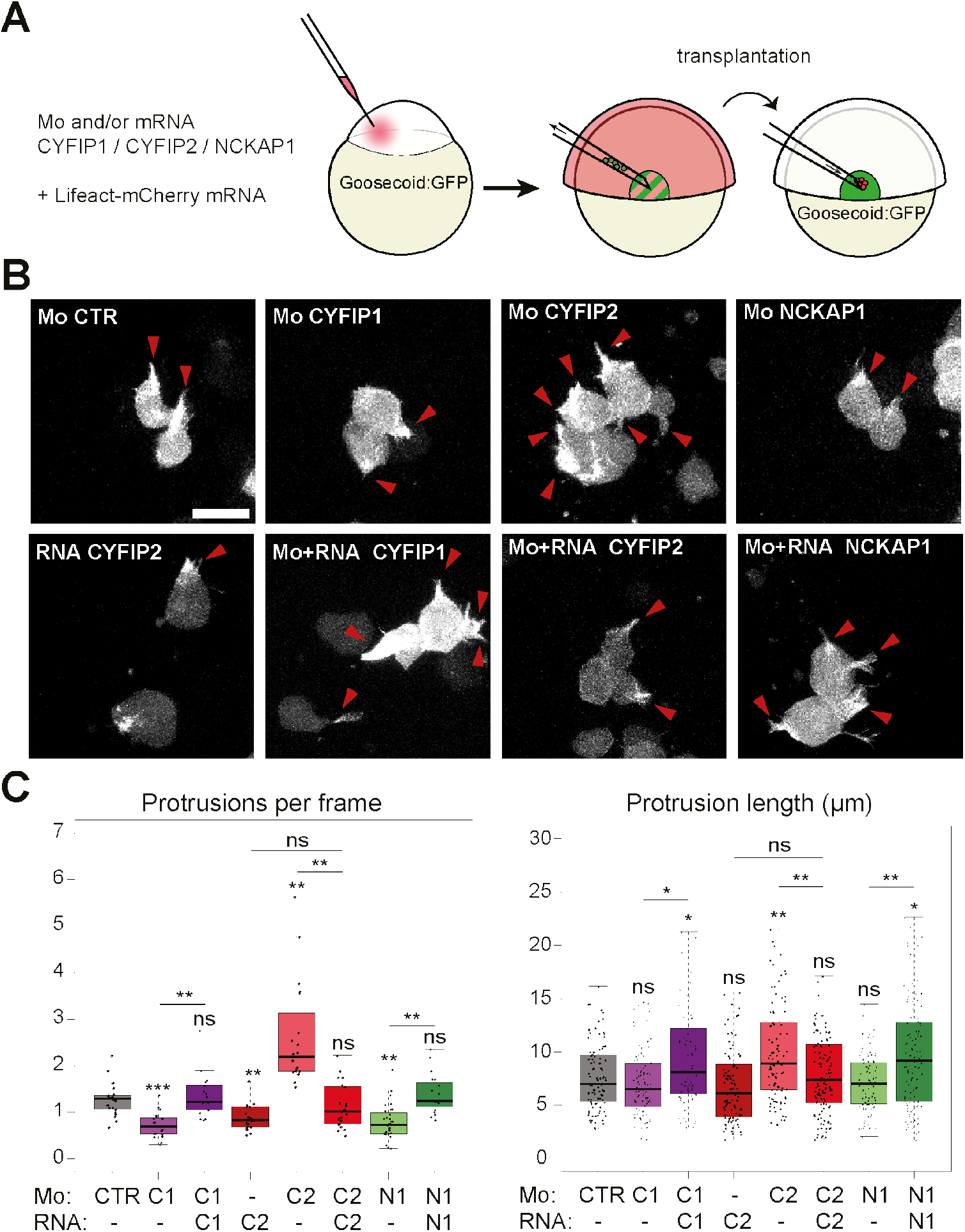
CYFIP2 depletion promotes actin rich protrusions in prechordal plate cells of zebrafish embryos. **A** Scheme of the experimental design. Donor embryos were injected with the actin filament marker Lifeact-mCherry mRNA and morpholinos (Mo) targeting a control sequence (CTR), CYFIP1 (C1), CYFIP2 (C2), NCKAP1 (N1), alone or in combination with mRNAs encoding the corresponding human proteins. Labeled prechordal plate cells from a donor embryo were transplanted into an uninjected embryo and recorded. **B** Images of representative cells with red arrowheads indicating actin-rich protrusions. Scale bar: 20 μm. **C** Box plot representations of protrusion numbers per frame (n=17 to 32 cells from 4 to 5 embryos per condition) and protrusion lengths (n=95 randomly selected protrusions per condition). Oneway ANOVA on linear mixed model accounting for the sampling biases. * P<0.05; ** P<0.01; *** P<0.001; ns: not significant. The p-values without a bar refer to comparisons with the control condition.

Together, the results of loss and gain-of-function experiments of CYFIP2 using zebrafish embryos are consistent with those obtained in human breast cells and demonstrate that the novel anti-migratory function of CYFIP2 is a conserved function of this subunit across vertebrates.

### CYFIP2 rescues lamellipodium formation in *CYFIP1/2* double KO cells

To examine whether CYFIP2 was a functional subunit of the WAVE complex, we re-expressed CYFIP2 in the published *CYFIP1/2* double knock-out B16-F1 cells (DKO) (Schaks *et al*, 2018). Similar to GFP-CYFIP1, GFP-CYFIP2 rescued lamellipodium formation in DKO cells, although CYFIP1 induced more prominent lamellipodia than CYFIP2 (Fig.6AB, Movie S6). The expression of CYFIP2 was unable to restore the full speed of protrusions observed in parental B16-F1 cells or in CYFIP1-rescued DKO cells (Fig.6B). Consistently, the branched actin machinery, composed of the filament branching Arp2/3 complex and cortactin that stabilizes Arp2/3 at the branched junction (Molinie & Gautreau, 2018b), was not as efficiently recruited at the leading edge by CYFIP2 as by CYFIP1 (Fig.6CD).

**Figure 6.**
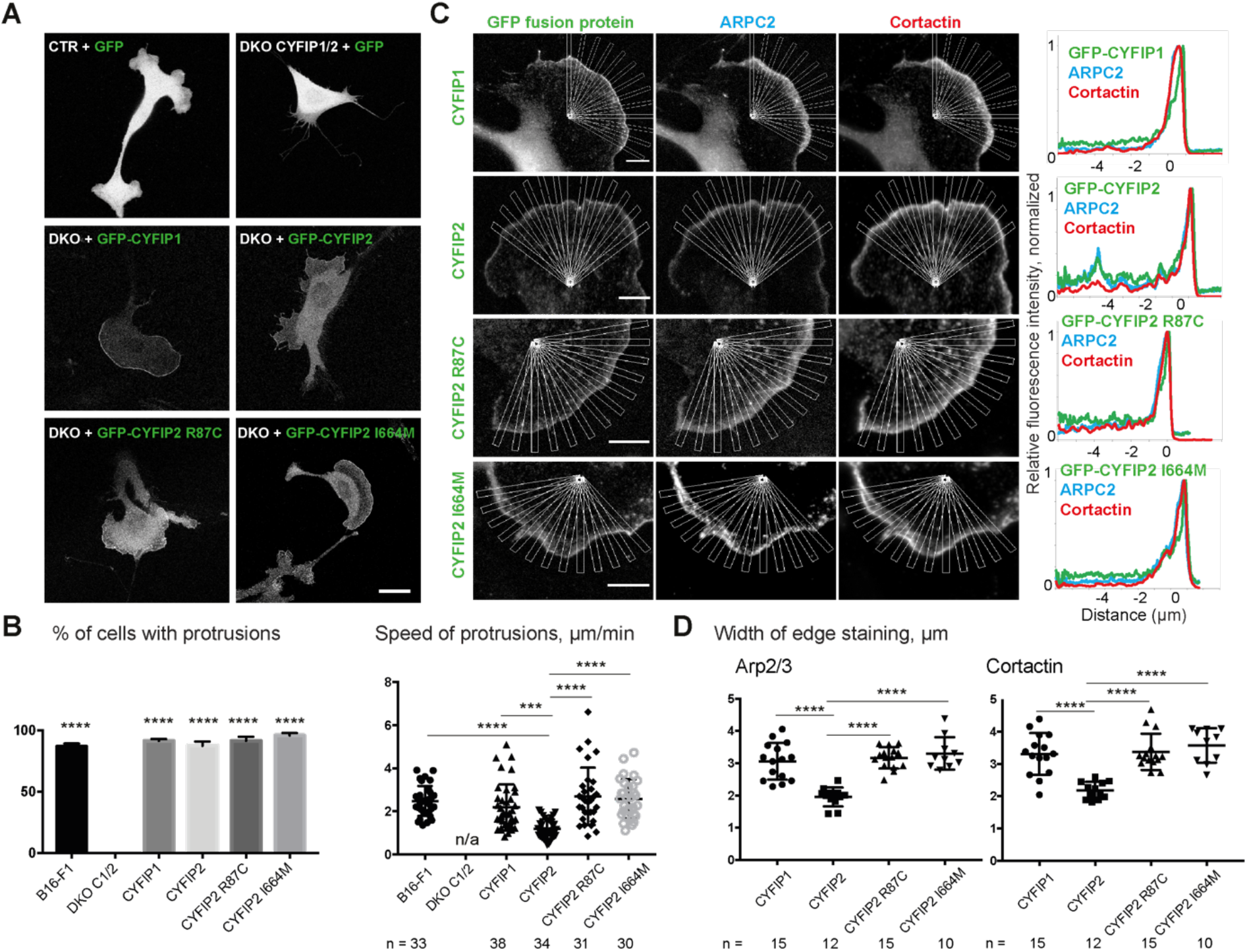
CYFIP2 rescues membrane protrusions of *CYFIP1/2* Double Knock-Out (DKO). **A** GFP-tagged CYFIP1, CYFIP2, and the two CYFIP2 mutant forms, R87C and I664M, were expressed in B16-F1 melanoma cells, control parental (CTR) or *CYFIP1/2* DKO cells as indicated. Distribution of GFP fusion proteins and morphology of transfected cells. Scale bar: 20 μm. **B** Percentage of transfected cells forming protrusions (n=100) and average speed of protrusions. Mean ± SD, one-way ANOVA. **C** Recruitment of GFP-tagged CYFIP1, CYFIP2, and mutant forms of CYFIP2 assessed by multiple radial line scans. Average profiles of the indicated markers upon registering line scans to the cell edge. Scale bars: 5μm. **D** Average width of Arp2/3 and cortactin staining. Mean ± SD. One-way ANOVA: ***P<0.001; ****P<0.0001.

We also analyzed the role of two point mutations of CYFIP2, R87C and I664M, which are recurring mutations found in patients affected by intellectual disability (Zweier *et al*, 2019). These two point mutations did not impair the ability of CYFIP2 to induce lamellipodia (Fig.6AB). On the opposite, mutated CYFIP2 seemed to induce more prominent lamellipodia than wild type (Movie S7). Mutant forms of CYFIP2 induced faster membrane protrusions than wild type, in line with increased recruitment of the branched actin machinery (Fig.6B-D). Mutant forms of CYFIP2 reached the activity of CYFIP1 in these assays. So CYFIP2 is a functional CYFIP protein, which is less active than CYFIP1, but whose activity can be enhanced by point mutations found in patients with neurodevelopmental defects.

### CYFIP2-containing WAVE complexes are less activatable by active RAC1 than CYFIP1-containing ones

Human CYFIP2 is 88 % identical to human CYFIP1. To understand the difference between CYFIP2 and CYFIP1-containing WAVE complexes, we built a homology model of the CYFIP2-containing complex by replacing CYFIP1 with CYFIP2 in the crystal structure of the WAVE complex, which contained WAVE1 lacking its central proline-rich region and ABI2 lacking its C-terminus (Chen *et al*, 2010). None of subunit-subunit interfaces were affected by the substitutions in CYFIP2 (Fig.7A). Consistently, all detectable CYFIP2 from a cytosolic lysate of MCF10A cells migrated at the position of the native WAVE complex by ultracentrifugation on sucrose gradients (Fig.7B)(Gautreau *et al*, 2004).

**Figure 7.**
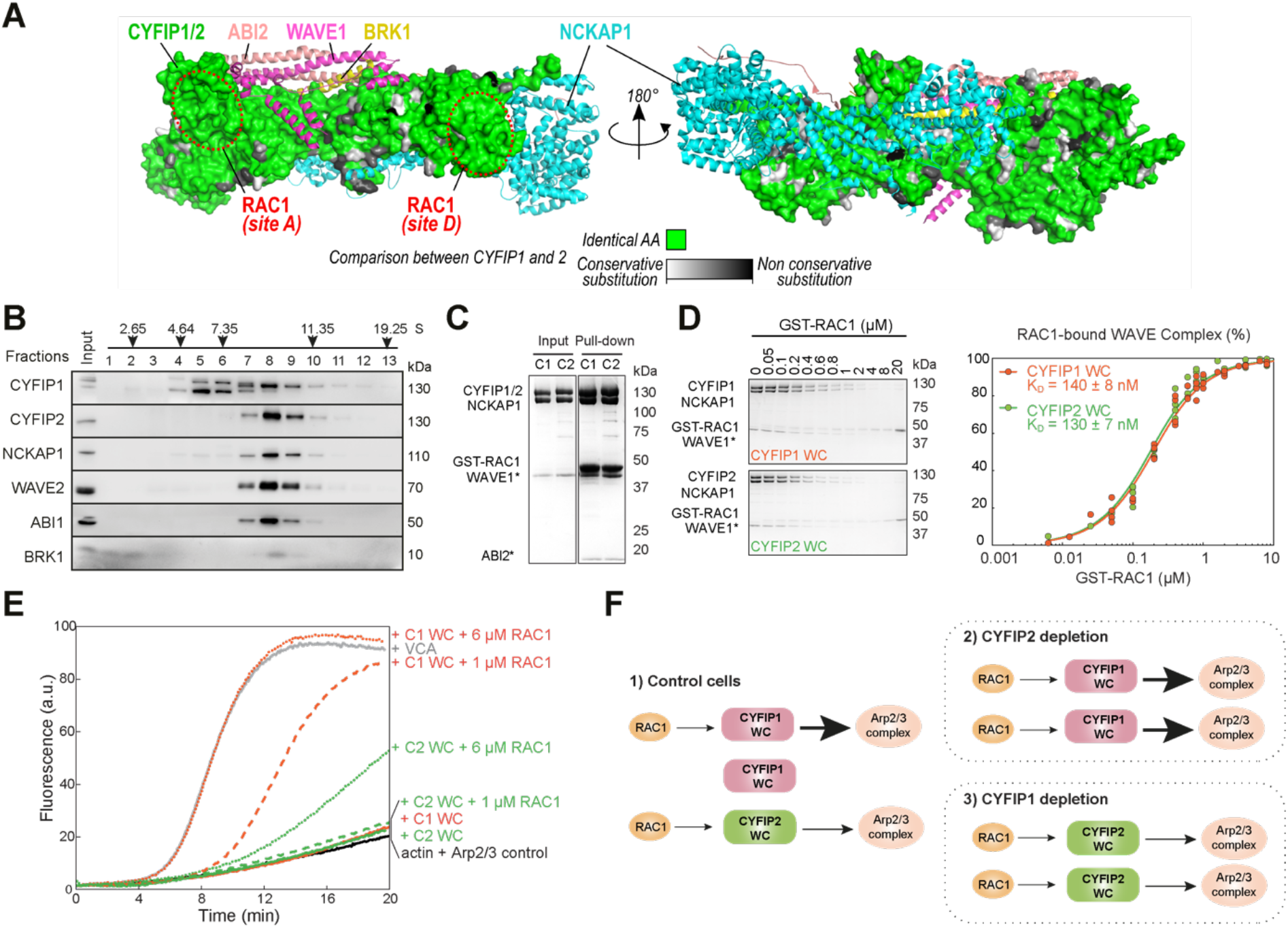
CYFIP2-containing WAVE complexes are less activatable by RAC1 than CYFIP1-containing WAVE complexes. **A** Structural models of CYFIP1 and CYFIP2. Sequence identity between CYFIP1 and CYFIP2 is 88% and non-conserved positions are color-coded. The vast majority of non-conserved residues fall outside of binding sites for known protein partners. WAVE complex subunits were obtained from PDB:4N78. **B** Ultracentrifugation of MCF10A cytosol on a sucrose gradient. WAVE complex subunits are revealed by Western blots. The CYFIP1 antibody cross-reacts with a lower molecular weight band. **C** Coomassie-blue stained SDS-PAGE gels showing reconstitution of WAVE complexes containing CYFIP1 or CYFIP2 and pull-down with GTP-bound RAC1 (GST-RAC1 Q61L P29S). ABI2* and WAVE1* are not full-length proteins. **D** Supernatants containing the WAVE complexes upon pull-down with increasing amounts of GST-RAC1 Q61L P29S. Dissociation constants K_D_ and standard errors are derived from fitting 4 independent experiments at various concentrations. **E** Pyrene-actin polymerization assay of CYFIP1- or CYFIP2-containing WAVE complexes. Conditions: 4 μM actin (5% pyrene-labeled), 10 nM Arp2/3 complex, 100 nM WAVE complexes (WC) or WAVE1 WCA, and indicated amounts of untagged RAC1 Q61L P29S. A single experiment is displayed, a second experiment gave similar results. **F** Model: CYFIP2-containing WAVE complexes activate less Arp2/3 upon RAC1 binding than CYFIP1-containing WAVE complexes. Upon depletion of CYFIP2, Arp2/3 activity increases, because more RAC1 is available to activate CYFIP1 containing complexes, leading to active membrane protrusions and increased migration persistence. On the opposite, upon depletion of CYFIP1, Arp2/3 activity decreases, because CYFIP2-containing complexes are poorly activated by RAC1, leading to weak membrane protrusions and decreased migration persistence. Increase in CYFIP2 levels upon CYFIP1 depletion does not compensate for the loss of CYFIP1-containing complexes.

Active RAC1 binds to two binding sites on the CYFIP1 subunit at the surface of the WAVE complex: the A site and the D site (Chen *et al*, 2017; Ding *et al*, 2022). Neither RAC1 binding sites were affected by substitutions in CYFIP2. We reconstituted a WAVE complex with either CYFIP1 or CYFIP2, using a previously described procedure (Chen *et al*, 2014), to examine its binding to and activation by RAC1. The two complexes indeed interacted equally well with GTP-bound RAC1 (Fig.7C and D). In pyrene-actin polymerization assays, however, the CYFIP2-containing WAVE complex was poorly activated by RAC1 compared to the CYFIP1-containing WAVE complex (Fig.7E). These in vitro data are thus consistent with the observation that in cells, CYFIP2 promotes membrane protrusions, but not as efficiently as CYFIP1.

## Discussion

CYFIP2 is highly related to CYFIP1. Both CYFIP proteins were previously reported to incorporate into WAVE complexes (Stradal *et al*, 2001; Kumar *et al*, 2013; Wan *et al*, 2015), a point we confirmed here through reconstitution of recombinant WAVE complexes and by analyzing endogenous complexes using sucrose gradients. Despite being a bona fide subunit of the WAVE complex that activates the Arp2/3 complex, CYFIP2 was found here to oppose cell migration in a variety of cell systems, MCF10A, MCF7, MDA-MB-231 and prechordal plate cells from the zebrafish embryo. In these experiments, CYFIP2 depletion enhanced cell migration, whereas CYFIP2 overexpression decreased cell migration.

Reconstituted complexes unambiguously showed that CYFIP2-containing WAVE complexes are poorly activated compared to CYFIP1-containing WAVE complexes, even though RAC1-GTP binds equally well to both complexes. Structural models of both complexes were predicting that RAC1 binding should not be affected, but did not allow to anticipate this poor activation of CYFIP2-containing WAVE complexes. This property accounts for the observed phenotypes. Indeed, depletion of CYFIP2 reallocates available GTP-bound RAC1 to readily activatable CYFIP1-containing WAVE complexes (Fig.7F). Overall, levels of branched actin at the cortex of a cell would not only be the result of RAC1 activation, but also of the balance between more or less activatable WAVE complexes. In line with this idea, reducing expression of CYFIP1 or CYFIP2 through gene dosage in heterozygous mice suffices to reveal neurobehavioral phenotypes (Babbs *et al*, 2019; Lee *et al*, 2020). These haploinsufficiencies highlight the importance of the balance between WAVE complexes containing CYFIP1 or CYFIP2.

Similar positive or negative modulations of actin polymerization were previously reported as a function of Arp2/3 composition for the two pairs of paralogous subunits, ARPC1A/ARPC1B and ARPC5/ARPC5L (Abella *et al*, 2016). Arp2/3 complexes containing ARPC1B and ARPC5L are more activatable by NPFs than the ones containing ARPC1A and ARPC5 and produce actin networks, whose branches are more stable. In line with these differential activities, depletion of ARPC1B decreases persistence of cell migration, whereas depletion of ARPC1A increases it (Molinie *et al*, 2019). For the WAVE complex, the VCA of WAVE1 significantly restrains elongation of actin filaments compared to the VCA of WAVE2 and that leads to an increased retrograde flow of actin from the lamellipodium tip when WAVE1 is depleted (Sweeney *et al*, 2015; Tang *et al*, 2020). This increased retrograde flow upon WAVE1 depletion does not, however, translate into faster membrane protrusions, nor increased cell migration. Unlike CYFIP2, the WAVE1 subunit that displays a lower activity than WAVE2 does not oppose cell migration. Mammalian genomes encoding paralogous subunits for many stable multiprotein complexes offer numerous opportunities to fine tune complex cellular responses, such as cell migration, through the co-expression of paralogous genes with different activities.

We found that the two recurring *CYFIP2* mutations found in children affected with intellectual disability and epileptic encephalopathy (Zweier *et al*, 2019), R87C and I664M, render CYFIP2 as active as CYFIP1 in restoring lamellipodia in *CYFIP1/2* DKO melanoma cells. It was previously reported that equivalent mutations when introduced into CYFIP1 also rendered the WAVE complex more active (Schaks *et al*, 2020). It was predicted and experimentally verified that CYFIP subunits harboring the R87C mutation assemble WAVE complexes that become independent from input RAC1 signaling because the output VCA domain would be less efficiently masked in presence of the R87C mutation (Nakashima *et al*, 2018; Schaks *et al*, 2020; Begemann *et al*, 2021). The severe neurological disorder due to such gain-of-function mutations emphasizes the importance for CYFIP2 to have the restrained activity we report here.

## Materials and Methods

### Cell lines, transfection and establishment of stable clones

MCF10A, MCF7 and MDA-MB-231 were from the collection of breast cell lines organized by Thierry Dubois (Institut Curie, Paris). 293 Flp-In was purchased from ThermoFisher Scientific. B16-F1 mouse melanoma cells that are *CYFIP1/2* double KO were a kind gift of Klemens Rottner (Helmholtz-Zentrum für Infektionsforschung, Braunschweig). MCF10A cells were grown in DMEM/F12 medium supplemented with 5% horse serum, 20 ng/mL epidermal growth factor, 10 μg/mL insulin, 500 ng/mL hydrocortisone, and 100 ng/mL cholera toxin. MDA-MB-231, MCF7, 293 Flp-In and B16-F1 cells were grown in DMEM medium with 10% FBS. Medium and supplements were from Life Technologies and Sigma. Cells were incubated at 37°C in 5% CO_2_.

Stable MCF10A cells expressing CYFIP2 were obtained by transfecting MCF10A cells, with the home-made plasmid MXS AAVS1L SA2A Puro bGHpA EF1Flag GFP CYFIP2 Sv40pA AAVS1R, or MXS AAVS1L SA2A Puro bGHpA EF1Flag GFP Blue Sv40pA AAVS1R as a control. Transfection was performed with Lipofectamine 2000 (Invitrogen). To obtain stable integration of the MXS plasmid at the AAVS1 site, cells were cotransfected with two TALEN plasmids inducing DNA double strand breaks at the AAVS1 locus (Addgene #59025 and 59026) (González *et al*, 2014). Cells were selected with 1 μg/mL puromycine (Invivogen) and pooled. Stable MCF10A cells expressing shRNA were obtained by transfection with previously described pSUPER-Retro-Puro plasmids (Steffen *et al*, 2004) and puromycin selection.

The stable 293 Flp-In cell line expressing Flag-HA-CYFIP1 was previously described (Derivery *et al*, 2009). An equivalent cell line expressing Flag-HA-CYFIP2 was obtained according to a published procedure (Derivery & Gautreau, 2010a).

MDA-MB-231, MCF7 and MCF10A were depleted by siRNAs (OnTarget Smart Pools, Dharmacon), transfected at 20 nM final concentration using Lipofectamine RNAiMAX (Invitrogen), and re-transfected 72h later, for a total of 6 days. For Fig.S4, siRNAs #1 and #2 of each pool were ordered and used.

The MCF10A CYFIP2 knockout cell line was generated with CRISPR/Cas9 system. The targeting sequence 5’-CAUUUGUCACGGGCAUUGCA-3’ was used to induce the double strand break. For the negative control the non-targeting sequence 5’-AAAUGUGAGAUCAGAGUAAU-3’ was used. Cells were transfected with crRNA:trackRNA duplex and the purified Cas9 protein by Lipofectamine CRISPRMAX™ Cas9 Transfection Reagent (all reagents from Thermofisher Scientific). The next day, cells were subjected to dilution at 0.8 cells/well in 96 well plates. Single clones were expanded and analyzed by CYFIP2 Western blot. From about 100 clones, 2 clones lacking CYFIP2 expression were identified. The PCR products amplified from genomic DNA containing the gRNA recognition site were then cloned (Zero Blunt PCR Cloning Kit, Thermofisher Scientific) and sequenced. Alleles were knock-out due to a frameshift of either +1 or −1 in the third exon of the *CYFIP2* gene in both clones.

### Antibodies and Western blot

Cells were lysed in RIPA buffer and analyzed by Western blot. SDS-PAGE was performed using NuPAGE 4-12% Bis-Tris and 3-8% Tris-Acetate gels (Life Technologies). Nitrocellulose membranes were developed with horseradish peroxidase (HRP) coupled antibodies (Sigma) and SuperSignal West Femto chemiluminescent substrate (Thermo Fisher Scientific). Homemade rabbit polyclonal antibodies against CYFIP1, ABI1, WAVE2 were previously described (Gautreau *et al*, 2004). The mouse monoclonal antibody, 231H9, targeting BRK1 was previously described (Derivery *et al*, 2008). The antibodies targeting CYFIP-2 (Sigma SAB2701081), NCKAP1 (Bethyl A305-178A), cortactin (Millipore 4F11), ARPC2 (Millipore 07-227) and tubulin (Sigma T9026) were purchased. Quantification of Western blots was performed by densitometry of boxed regions of interest with background subtraction, using the ImageJ software. Densitometry values were converted into ng amount when and only when the value felt within the range of an internal standard curve of purified CYFIP1/2 containing WAVE complexes and that the standard curve gave linear results.

### Sucrose gradient

For sucrose gradient analysis of WAVE subunits, Nitrogen cavitation (Parr instruments, 500 Psi for 20 min) followed by centrifugation (16,000 × g, 20 min) and ultracentrifugation (150,000 × g, 60 min) were used to prepare cytosolic extracts from cells trypsinized from two 15 cm dishes and resuspended in the XB buffer (20 mM HEPES, 100 mM NaCl, 1mM MgCl_2_, 0.1 mM EDTA, 1mM DTT, pH 7.7). 200 μL of extract was loaded on the 11 mL 5–20% sucrose gradient in the XB buffer and subjected to ultracentrifugation for 17 h at 197,000 ×g in the swinging bucket rotor SW41 Ti (Beckman). 0.5 mL fractions were collected and concentrated by using trichloroacetic acid precipitation with insulin as a carrier. The samples were washed with acetone, dried and then resuspended in the 1x LDS loading buffer with 2.5% of β-ME for Western blot analysis.

### Migration assays

Transwell migration assays were performed using FluoroBlok inserts with 8 μm holes (Corning, 351152), covered with 20 μg/ml fibronectin (Sigma, F1141). MDA-MB-231 cells were plated in serum-free medium and allowed to migrate towards serum-containing medium for 16 h, incubated with 4 μg/ml calcein AM (Sigma, C1359) for 1 h, and images of fluorescent cells were acquired and quantified using ImageJ software.

Assays of random single cell migration in 2D were performed in 8 chamber Ibidi dishes (Biovalley 80826) covered with 20 μg/ml fibronectin. Assays of wound healing were performed by lifting an Ibidi insert as previously described (Molinie & Gautreau, 2018a). 3D migration was performed in 2 mg/ml collagen gel polymerized at 37°C (rat tail collagen type I, Corning 354263), with the cells sandwiched between the two layers of collagen. An inverted Axio Observer microscope (Zeiss) equipped with a Pecon Zeiss incubator XL multi S1 RED LS (Heating Unit XL S, Temp module, CO_2_ module, Heating Insert PS and CO_2_ cover), a definite focus module and a Hamamatsu camera C10600 Orca-R2 was used to perform videomicroscopy. Pictures were taken every 5 min for 24 h for 2D migration, and every 20 min for 48 h for 3D migration. Migration parameters including migration persistence, based on the angular shift of displacement vectors between frames, was analyzed as previously described (Dang *et al*, 2013) using DiPer programs (Gorelik & Gautreau, 2014).

### Imaging protrusions

GFP-tagged human CYFIP1 or CYFIP2 (wild type or mutant) was transiently transfected into the B16 double KO cells, and 48 h later, 10-minute videos (images taken every 10 seconds) were acquired using a confocal laser scanning microscope (TCS SP8, Leica) equipped with a high NA oil immersion objective (HC PL APO 63×/ 1.40, Leica), a white light laser (WLL, Leica) and controlled by the LasX software. Protrusion speed was measured using the Multi Kymograph tool in ImageJ software. For the LineScan analysis, images of fixed, stained cells were obtained, and analyzed as described in (Dang *et al*, 2013) and (Molinie *et al*, 2019).

### Zebrafish embryos, cell transplantation and imaging

Embryos were obtained by natural spawning of *Tg(−1.8gsc:GFP)ml1* adult fishes (Doitsidou *et al*, 2002). All animal studies were done in accordance with the guidelines issued by the Ministère de l’Education Nationale, de l’Enseignement Supérieur et de la Recherche and were approved by the Direction Départementale des Services Vétérinaires de l’Essonne and the Ethical Committee N°59.

Translation blocking morpholinos (Gene Tool LLC Philomath) were designed against zebrafish *CYFIP1* (AAAAACTATCCGCTTCGACTGTTCA) and *CYFIP2* (CGACACAGGTTCACTCACAAAACAG). The *NCKAP1* morpholino (CCGAGACATGGCTCAAACGACCGTC) was described in (Biswas *et al*, 2010). The control morpholino is a standard control (CCTCTTACCTCAGTTACAATTTATA). mRNAs were synthesized using pCS2+ plasmids containing the human genes described in (Gautreau *et al*,2004) and the mMessage mMachine SP6 kit (Thermo Fischer).

For cell migration quantification, embryos were injected at the one-cell stage with 1.5 nl of a solution containing Histone2B-mCherry mRNA (30 ng/μl) and either control morpholino (0.1, 0.2 or 0.8mM), MoCYFIP1 (0.2mM), MoCYFIP2 (0.1mM) or MoNCKAP1 (0.8mM), with or without mRNAs encoding either human CYFIP1 (10ng/μl), human CYFIP2 (10ng/μl) or human NCKAP1 (10ng/μl). Injected embryos were mounted in 0.2% agarose in embryo medium and imaged between 60% and 80% epiboly (6.5-8.5 hpf) under an upright TriM Scope II (La Vision Biotech) two photon microscope equipped with an environmental chamber (okolab) at 28°C using a 25x water immersion objective. Visualization of 3D movies and nuclei tracking were done using Imaris (Bitplane). Cell migration parameters were extracted using custom Matlab (Math Works) code and autocorrelation was computed using published Excel macros (Gorelik & Gautreau, 2014).

For protrusion analysis, embryos were injected in one cell at the four-cell stage with 1.5 nl of a solution containing Lifeact-mCherry mRNA (50 ng/μl) and either control morpholino (0.5 mM), MoCYFIP1 (0.2mM), MoCYFIP2 (0.1mM) or MoNCKAP1 (0.8mM), with or without mRNAs encoding either human CYFIP1 (10 ng/μl), human CYFIP2 (10 ng/μl) or human NCKAP1 (10 ng/μl). Small cell groups were transplanted at shield stage (6 hpf) from the shield of an injected embryo to the shield of an untreated host. Embryos were then cultured in embryo medium (Hans *et al*, 2007) with 10 U/mL penicillin and 10 μg/mL streptomycin. Transplanted embryos were mounted in 0.2% agarose in embryo medium and imaged between 60% and 80% epiboly (6.5-8.5 hpf) under an inverted TCS SP8 confocal microscope equipped with environmental chamber (Leica) at 28°C using a HC PL APO 40x/1.10 W CS2 objective. Visualization of images was done on ImageJ, lamellipodia-like actin rich protrusions being quantified on the basis of morphological criteria as described in (Diz-Muñoz *et al*, 2010).

### Reconstitution of WAVE complexes and in vitro assays

Recombinant WAVE complexes containing full-length human CYFIP1 or CYFIP2, full-length NCKAP1, full-length BRK1, ABI2 (1-158) and WAVE1 (1-230)-(GGS)_6_-WCA (485-559), referred to as WRC230WCA were purified as previously described (Chen *et al*, 2014, 2017). CYFIP1-and CYFIP2-containing WAVE complexes behaved similarly during expression and purification by various chromatographic steps. Other proteins, including the Arp2/3 complex, actin, WAVE1 WCA, Tev, GST-RAC1 (Q61L P29S, 1-188), and untagged RAC1 (Q61L P29S, 1-188) were purified as previously described (Chen *et al*, 2017).

GST pull-down experiments were performed as previously described (Chen *et al*, 2017). Briefly, 200 pmol of GST-RAC1 and 200 pmol of WAVE complex were mixed with 20 μL of Glutathione Sepharose beads (GE Healthcare) in 1 mL of binding buffer (10 mM HEPES pH 7, 50 or 100 mM NaCl, 5% (w/v) glycerol, 2 mM MgCl_2_, 1 mM DTT, and 0.05% Triton X100) at 4 °C for 30 min, followed by three washes using 1 mL of the binding buffer in each wash. Finally, the bound proteins were eluted with GST elution buffer (100 mM Tris-HCl pH 8.5, 30 mM reduced glutathione, and 2 mM MgCl_2_) and examined on SDS-PAGE gels.

GST equilibrium pull-down assays were performed in the EPD buffer (10 mM HEPES pH 7, 50 mM NaCl, 5% (w/v) glycerol, 2 mM MgCl_2_, and 1 mM DTT) as previous described (Chen et al., 2017). Essentially, each 100 μL of reaction contained 0.1 μM WRC230WCA, varying concentrations of GST-Rac1(Q61L P29S, 1-188), 30 μL of the Glutathione Sepharose beads, and 0.05% Triton X100. All protein samples and beads were first dialyzed or equilibrated in the EPD buffer prior to use. After gentle mixing at 4°C for 30 min, the beads were pelleted by a brief centrifugation, and the supernatant was immediately transferred to SDS loading buffer and analyzed by Coomassie blue-stained SDS-PAGE gels. Total intensity of the CYFIP1/2 and NCKAP1 bands was quantified by ImageJ to determine the unbound WAVE complex. The derived fractional occupancy from several independent experiments was pooled and globally fitted to obtain the binding isotherms and the apparent dissociation constants K_D_.

Actin polymerization assays were performed as previously described (Chen *et al*, 2017) with slight modifications. Each reaction (120 μL) contained 4 μM actin (5% pyrene labeled), 10 nM Arp2/3 complex, 100 nM WRC230WCA or WAVE1 WCA, and desired concentration of untagged RAC1 (Q61L P29S, 1-188) in NMEH20GD buffer (50 mM NaCl, 1 mM MgCl_2_, 1 mM EGTA, 10 mM HEPES pH7.0, 20% (w/v) glycerol, and 1 mM DTT). Pyrene-actin fluorescence was recorded every 5 seconds at 22 °C using a 96-well flat-bottom black plate (Greiner Bio-One™) in a Spark plater reader (Tecan), with excitation at 365 nm and emission at 407 nm (15 nm bandwidth for both wavelengths).

### Statistical analyses

Exponential decay and plateau fit of migration persistence 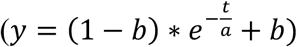 were performed for all individual cells. Statistical analysis was performed in R using linear mixed-effect models to take into account the resampling of the same statistical unit. Coefficients were then compared using one-way ANOVA. Multiple comparisons were performed using post hoc tests, Tukey’s for figure 6 and Dunnett’s for all others. Differences were considered significant at confidence levels greater than 95% (p < 0.05). Four levels of statistical significance were distinguished: *P<0.05; **P<0.01; ***P<0.001; ****P<0.0001.

## Acknowledgements

We thank Nelia Cordeiro for technical support, Theresia Stradal for sharing shRNA plasmids targeting NCKAP1 and CYFIP2, Klemens Rottner for the *CYFIP1/2* double KO cells, and Chuang Yu for help with statistics. This work was supported by grants from the Agence Nationale de la Recherche (ANR-20-CE13-0016-01 for AMG and NBD) and from National Institute of Health (grant R35 GM128786) to BC. We thank the Polytechnique Bioimaging Facility for assistance with live imaging on their equipment partly supported by Région Ile-de-France (interDIM) and Agence Nationale de la Recherche (ANR-11-EQPX-0029 Morphoscope2, ANR-10-INBS-04 France BioImaging).

## Author contributions

AP performed *in vitro* experiments of cell migration. AB and NBD performed *in vivo* experiments in zebrafish embryos. SY, YL and BC reconstituted the two CYFIP1/2 containing complexes and performed RAC1 binding/activation assays. RG performed the structural comparison of CYFIP1/2 proteins. SR performed the confocal microscopy of B16-F1 cells and helped with the data analysis. ML performed statistical modeling of migration persistence. YW performed the sucrose gradient experiments. NM constructed the integrative plasmid used to overexpress CYFIP2. AF and YW isolated the CYFIP2 knockout clones and performed experiments with these clones. NR quantified CYFIP1/2 protein expression in cell lines. AMG supervised the study. AP and AMG wrote the manuscript. All authors have commented on the manuscript and approved the submission.

