## Supplementary material for "Restrained activation of CYFIP2-containing WAVE complexes controls membrane protrusions and cell migration": Supp information

**Supplementary Information**

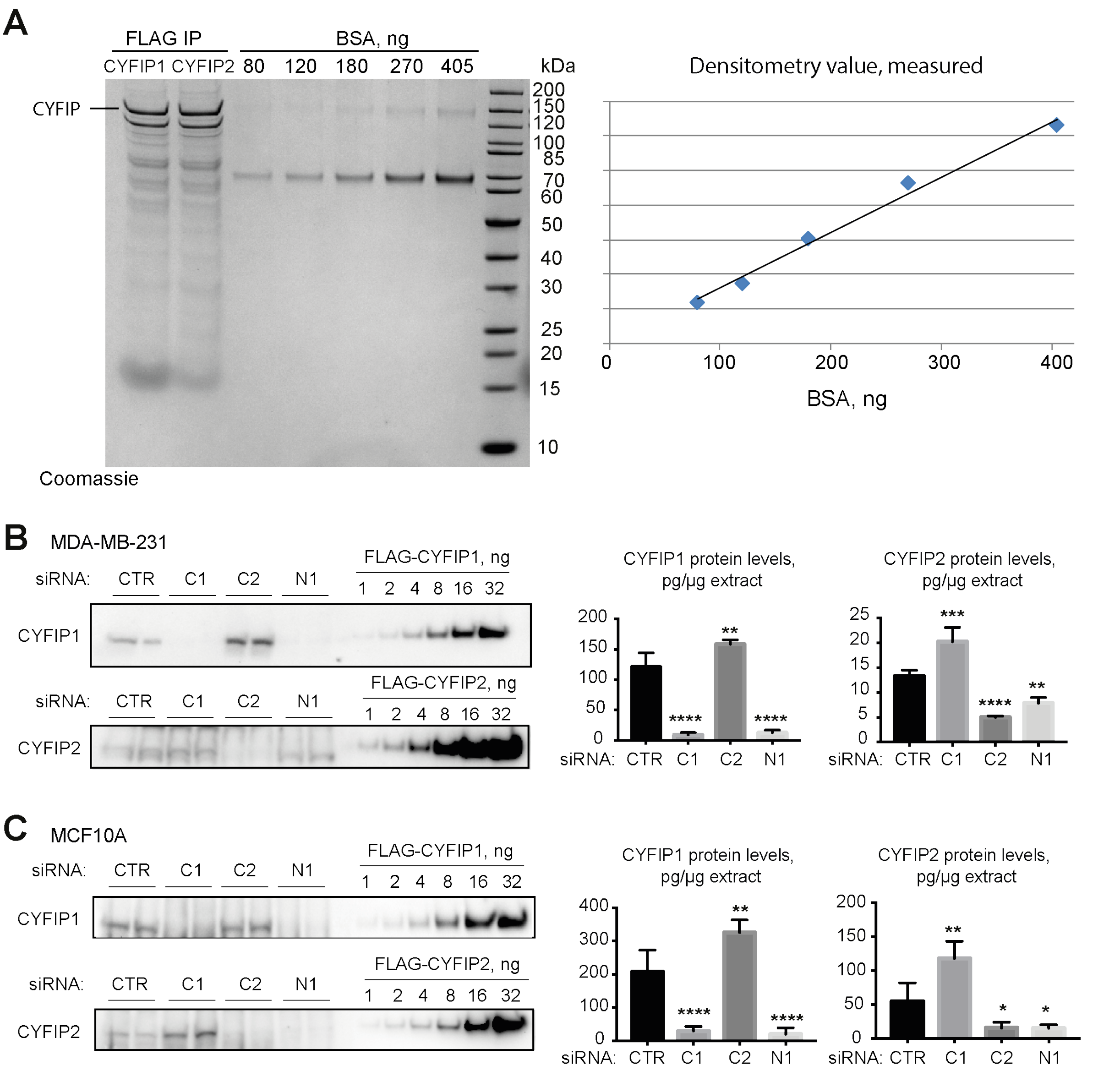


**Figure S1. Quantification of CYFIP1 and CYFIP2 proteins in MDA-MB-231 and MCF10A cell lysates. A** Coomassie-stained gel with FLAG-tagged immunopurified (IP) CYFIP1/2-containing WAVE complexes, and BSA standard curve. CYFIP1 or CYFIP2-containing WAVE complexes were purified from stable 293 Flp-In cell lines. **B** Quantification of CYFIP1 and CYFIP2 levels in RIPA extracts obtained from MDA-MB-231 cells transfected with the indicated siRNAs. Duplicate transfections of the siRNA smartpool were analyzed for each gene. The experiment was repeated twice (total n=4). **C** Exact same experiments performed with MCF10A cells. Mean ± SD. One-way ANOVA: * P<0.05; ** P<0.01; *** P<0.001; **** P<0.0001.

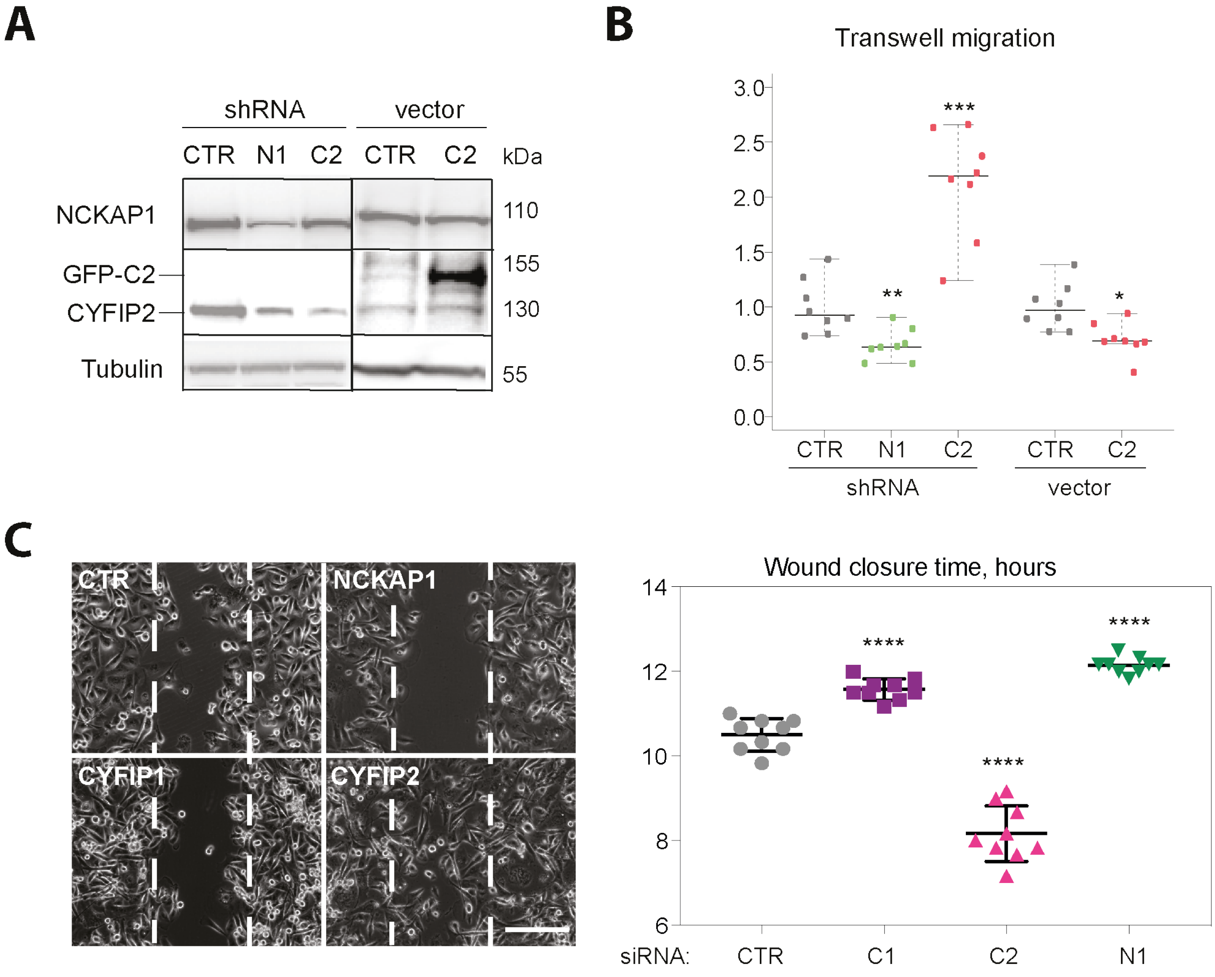


**Figure S2. Transwell migration and wound healing assays of MDA-MB-231 cells. A** stable MDA-MB-231 cell lines expressing either the indicated shRNAs or overexpressing the GFP-CYFIP2 protein (GFP-C2) were analyzed by Western blots with NCKAP1 and CYFIP2 antibodies. **B** Quantification of migration efficiency through transwell filters, mean ± SD of 9 measurements (3 biological repeats in triplicate), one-way ANOVA. **C** MDA-MB-231 cells were transfected with pools of siRNAs targeting CYFIP1 (C1), CYFIP2 (C2), NCKAP1 (N1) or non-targeting ones (CTR). Still images were extracted from movies at the time of the first complete closure. Scale bar, 400 µm. Quantification of the closure time, mean ± SD of 9 measurements (3 biological repeats in triplicate), one-way ANOVA. * P<0.05; *** P<0.001; **** P<0.0001.

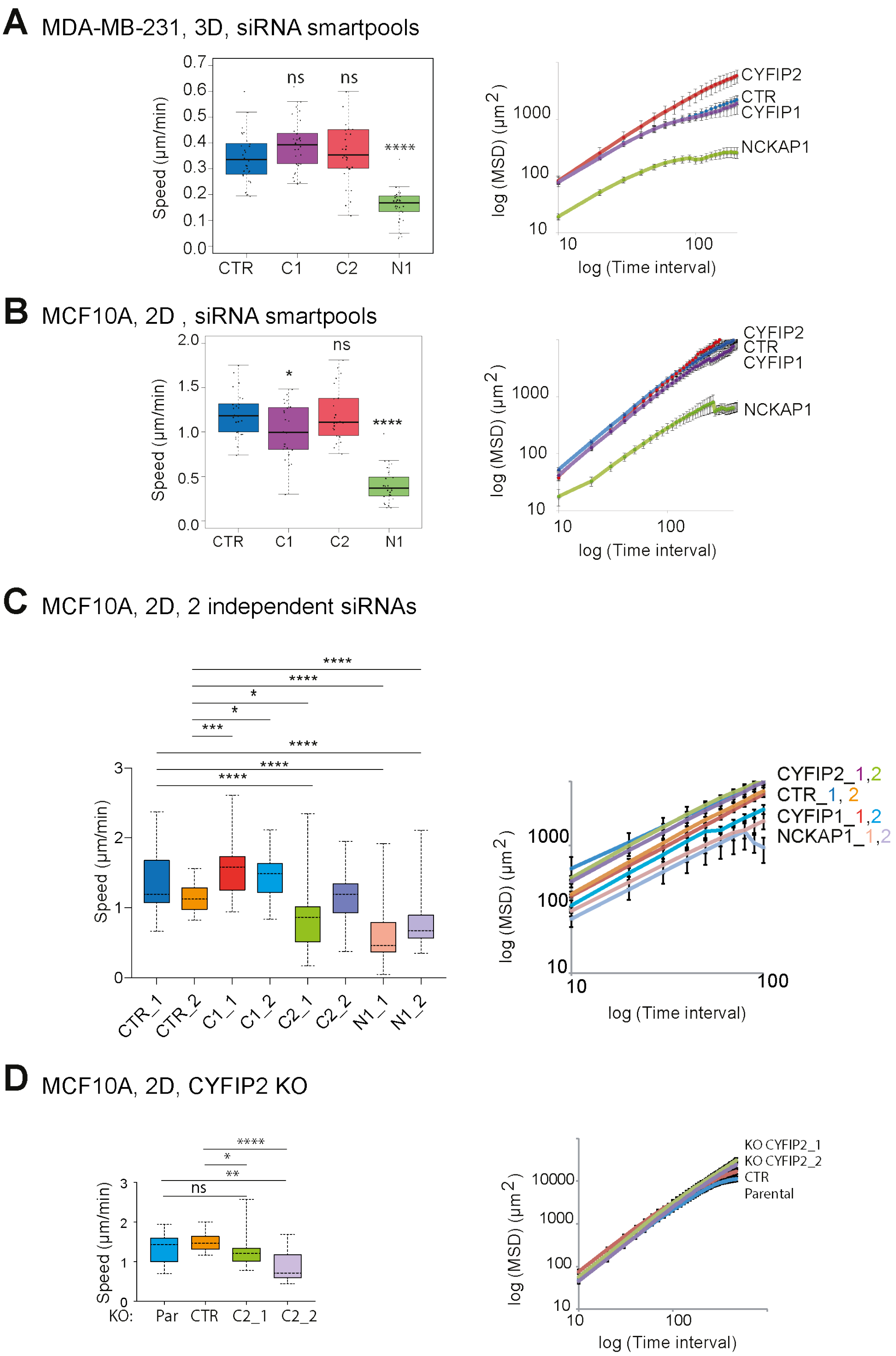


**Figure S3. Speed and MSD corresponding to cell trajectories.** **A** Experiments with MDA-MB-231 in 3D displayed in Fig.1D. n=30. **B** Experiments with MCF10A in 2D displayed in Fig.3D. n=25. **C** Experiments with MCF10A in 2D displayed in Fig.S4. n=30. **D** Experiments with MCF10A KO cells in 2D displayed in Fig.S5. n=30. One-way ANOVA: * $P<0.05$; ** $P<0.01$; *** $P<0.001$; **** $P<0.0001$; ns not significant.

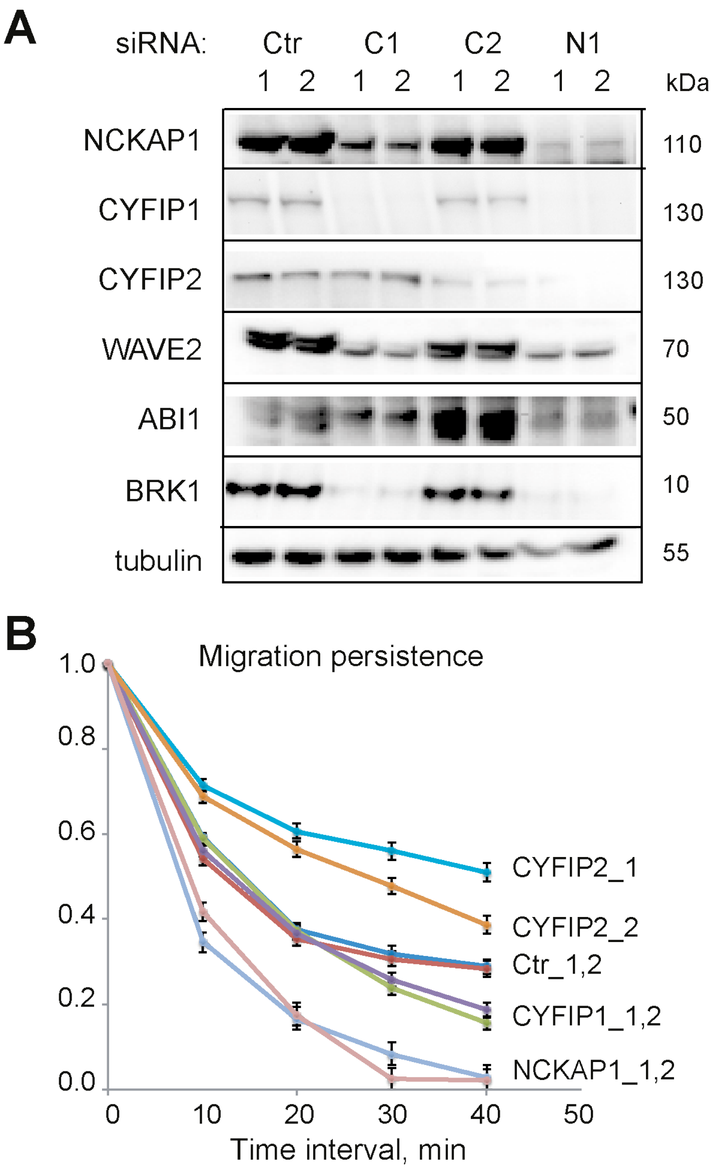


| | comparison | p-value | |
|---|---|---|---|
| 1 | Ctr_1/C1_1 | 0.46 | ns |
| 2 | Ctr_2/C1_1 | 0.82 | ns |
| 3 | Ctr_1/C1_2 | 0.38 | ns |
| 4 | Ctr_2/C1_2 | 0.71 | ns |
| 5 | Ctr_1/C2_1 | 0.0012 | ** |
| 6 | Ctr_2/C2_1 | 0.00023 | *** |
| 7 | Ctr_1/C2_2 | 0.044 | * |
| 8 | Ctr_2/C2_2 | 0.014 | * |
| 9 | Ctr_1/N1_1 | 0.00020 | *** |
| 10 | Ctr_2/N1_1 | 0.00077 | *** |
| 11 | Ctr_1/N1_2 | 0.00049 | *** |
| 12 | Ctr_2/N1_2 | 0.0016 | ** |

**Figure S4.** Two independent siRNAs targeting *CYFIP1*, *CYFIP2*, and *NCKAP1* were used to deplete the corresponding proteins from MCF10A cells. **A** Depletion was evaluated by Western blot. **B** From single cell trajectories of 2D random migration assays, migration persistence was calculated and plotted. n=30. One-way ANOVA.

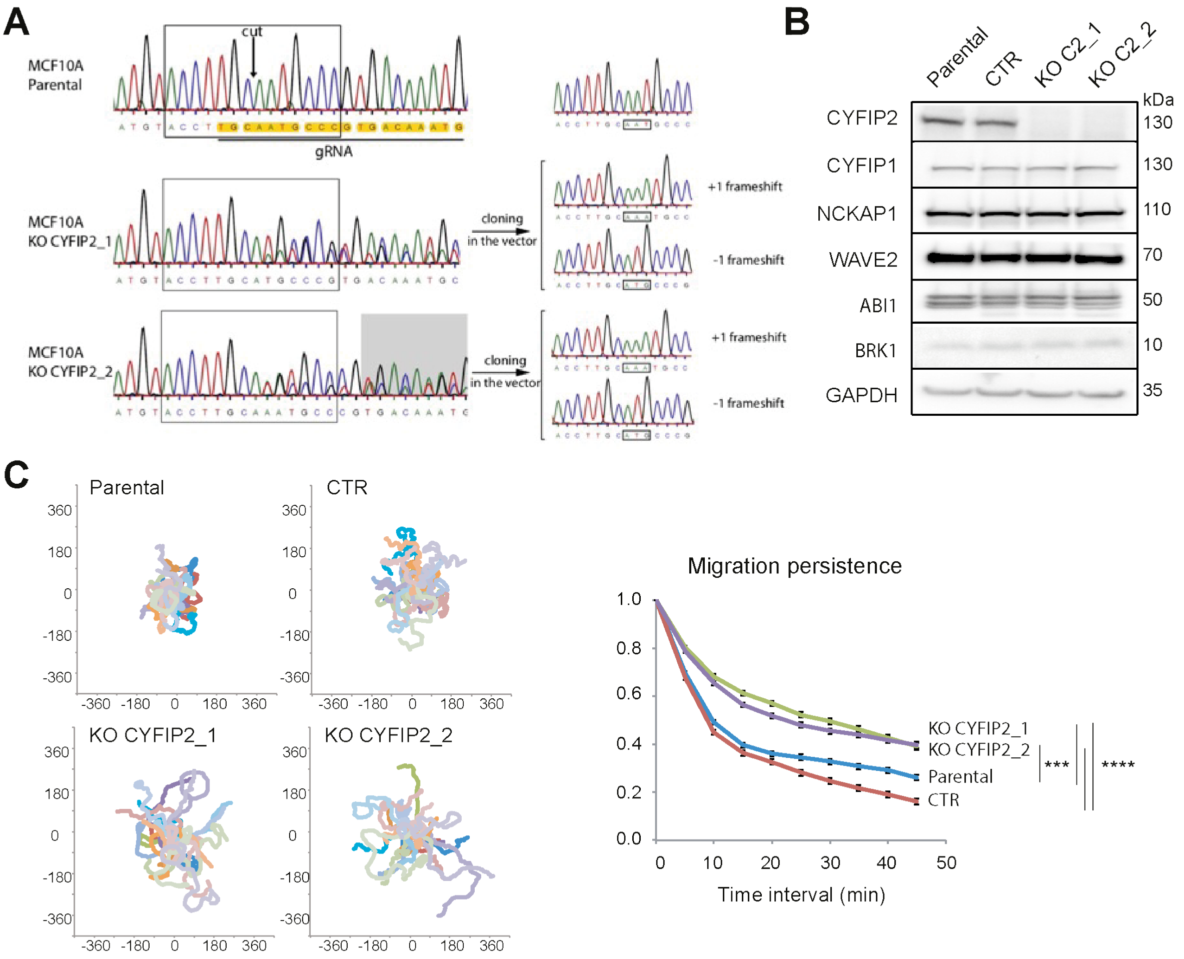


**Figure S5. Characterization of *CYFIP2* KO MCF10A clones**. **A** Genetic characterization of the two independent KO clones that were isolated. **B** Expression of WAVE complex subunits. **C** Trajectories of *CYFIP2* KO and control cells in 2D random migration assay. Migration persistence, n=30. One-way ANOVA: *** P<0.001; **** P<0.0001.

**Legends of movies**

**Movie S1. CYFIP2 depletion enhances membrane protrusions of MDA-MB-231 cells in 3D collagen gel.** Cells transfected with the indicated siRNAs targeting NCKAP1, CYFIP1, CYFIP2 or non-targeting controls (CTR) were recorded by phase contrast optics for 48 h with one frame every 20 min. Scale bar: 50 µm.

**Movie S2. CYFIP2 depletion promotes wound healing of MCF7 cells.** Cells were transfected with indicated siRNAs targeting NCKAP1, CYFIP1, CYFIP2 or non-targeting controls (CTR). A wound was obtained by lifting inserts and wound healing was monitored by videomicroscopy. A phase contrast image was taken every 20 min. Scale bar: 400 µm.

**Movie S3. CYFIP2 depletion promotes membrane protrusions and spreading of MCF10A cells.** Cells transfected with the indicated siRNAs targeting NCKAP1, CYFIP1, CYFIP2 or non-targeting controls (CTR) were recorded by phase contrast optics for 24 h with one frame every 5 min. For the calculation of migration parameters, only single cells were analyzed. Scale bar: 50 µm.

**Movie S4. Tracking of prechordal plate cells.** Nuclei of a Tg(gsc:GFP) embryo were labeled with Histone2B-mCherry (magenta). A Z-stack was acquired every 2 min. Nuclei of prechordal plate cells (identified by GFP expression and morphological criterion), not visible here, were 3D-tracked in time (white squares). Tracks are building up as cells are moving. Animal pole is to the right.

**Movie S5. CYFIP2 depletion promotes actin rich protrusions in prechordal plate cells.** Donor embryos were injected with the actin filament marker Lifeact-mCherry mRNA and morpholinos (Mo) targeting CYFIP1 (C1), CYFIP2 (C2), NCKAP1 (N1), alone or in combination with mRNAs encoding the corresponding human proteins. Red fluorescent prechordal plate cells were transplanted into a receiver embryo and Lifeact-mCherry fluorescence was imaged at 1 frame per minute. Scale bar: 25 µm.

**Movie S6. GFP-CYFIP2 restores lamellipodium protrusion and is recruited to the lamellipodial edge.** Control and *CYFIP1/2* double knock-out (DKO) B16-F1 cells were transfected with the indicated GFP plasmids and green fluorescence was recorded every 10 s for 10 min. Scale bar: 20 µm.

**Movie S7. R87C and I664M CYFIP2 restore lamellipodium protrusion and are recruited to the lamellipodial edge.** *CYFIP1/2* double knock-out (DKO) B16-F1 cells were transfected with the indicated GFP plasmids and green fluorescence was recorded every 10 s for 10 min. Scale bar: 20 µm.
